# High Bacterial Load Predicts Poor Outcomes in Patients with Idiopathic Pulmonary Fibrosis

**DOI:** 10.1101/003145

**Authors:** P L Molyneaux, M J Cox, S A G Willis-Owen, K E Russell, P Mallia, A M Russell, S L Johnston, A U Wells, W O C Cookson, T M Maher, M F Moffatt

**Author notes:** Corresponding author: Professor William O. C. Cookson National Heart and Lung Institute, Imperial College London, Guy Scadding Building, Royal Brompton Campus, London, SW3 6LY, UK. These three senior authors contributed equally to the study (W O C Cookson, T M Maher, M F Moffatt).

## Abstract

**Background:** Repetitive alveolar damage and aberrant repair may be important in the development of the fatal condition Idiopathic Pulmonary Fibrosis (IPF). The role played by microorganisms in this cycle is unknown.

**Methods:** We consecutively enrolled patients diagnosed with IPF according to international criteria together with healthy smokers, non-smokers and subjects with moderate Chronic Obstructive Pulmonary Disease (COPD) as controls. Subjects underwent bronchoalveolar lavage (BAL) from which genomic DNA was isolated. The V3-V5 region of the bacterial 16S rRNA gene was amplified, allowing quantification of bacterial load and identification of communities by 16S rRNA qPCR and pyrosequencing.

**Results:** Our 65 IPF patients had 3.9×10^9^ copies of the 16S rRNA gene per ml of BAL, two-fold more than the 1.8×10^9^ copies in 44 sex- and smoking-matched controls (*P*<0.0001). Baseline BAL bacterial burden predicted Forced Vital Capacity (FVC) decline (*P*=0.02). Patients in the highest tertile of bacterial burden were at a higher risk of mortality compared to subjects in the lowest tertile (hazard ratio 4.59 (95% CI, 1.05-20); *P*=0.04).

Sequencing yielded 912,883 high quality reads from all subjects. Operational Taxonomic Units (OTUs) representing *Haemophilus, Streptococcus, Neisseria* and *Veillonella* were 1.5 to 3.5 fold more abundant in cases than controls (*P*<0.05). Regression analyses indicated that these specific OTUs as well as bacterial burden associated independently with IPF.

**Conclusions:** IPF is characterised by an increased bacterial burden in BAL that predicts decline in lung function and death. Clinical trials of antimicrobial therapy may determine if microbial burden is causal or not in IPF progression.

## Introduction

Idiopathic pulmonary fibrosis (IPF) is a progressive and invariably fatal disease of unknown cause.^1^ It is increasing in prevalence^2^ and carries a similar mortality to lung cancer.^3^ Consistent clues to its aetiology have been that it occurs primarily in older adults, many of whom have been smokers, and that polymorphisms in genes related to epithelial integrity and host defence predispose to the disease.^4–8^ Epidemiological studies have highlighted occupational and domestic exposures associated with an increased risk of developing IPF,^9^ suggesting that environmental triggers may be integral to the pathogenesis of IPF in genetically susceptible individuals.

The histological pattern of fibrosis in IPF is consistent with multiple discrete alveolar epithelial injuries, supporting a model of repeated exposure and injury. Although factors which initiate the fibrotic process have not yet been identified, susceptibility to infection may be an important contributor to disease progression.^10^ While viruses may play a part in the initiation and progression of disease and also be responsible for a proportion of acute exacerbations,^11^ the role of bacteria in the pathogenesis and progression of IPF has not yet been studied in detail. Active infection in IPF is, however, known to carry a high morbidity and mortality.^12^ In individuals with IPF immunosuppression is clearly deleterious^13^ while treatment adherent subjects in a large trial of the use of prophylactic co-trimoxazole in IPF, experienced a reduction in overt infections and mortality.^14^

We therefore investigated the role of bacteria in the pathogenesis and progression of IPF in a substantial prospective case-control study with strict use of American Thoracic Society diagnostic criteria for the disease.^15^ We serially recruited and investigated patients at the time of first presentation, so that the subjects were representative of the disease spectrum attending our clinics. As IPF often appears against a background of smoking related lung disease, we included patients with COPD and matched numbers of smokers in our control groups. We used sequence-based culture-independent methodologies to quantify the numbers of bacterial genomes as well as the community composition of specimens collected by bronchoalveolar lavage.

## Methods

### Study Design

Patients undergoing diagnostic bronchoscopy with bronchoalveolar lavage (BAL) for suspected but previously undiagnosed IPF were consecutively recruited from the Interstitial Lung Disease Unit at the Royal Brompton Hospital, London, England between November 2010 and January 2013. In all cases bronchoscopy was undertaken at initial evaluation. Written informed consent was obtained from all subjects and the study was approved by the Local Research Ethics Committee (Ref 10/H0720/12). A diagnosis of IPF was made following multidisciplinary team discussion and individuals fulfilling currently accepted international criteria were included in the study.^15^ Subjects were excluded if they had a history of self-reported upper or lower respiratory tract infection, IPF acute exacerbation or antibiotic use in the prior three months. Subjects were regularly reviewed and lung function was repeated every six months.

Control subjects, including healthy individuals (smokers and non-smokers) and individuals with moderate (GOLD stage II) Chronic Obstructive Pulmonary Disease (COPD) were recruited separately using the same protocols. Recruitment of controls began before and continued during recruitment of IPF subjects. Subjects with other respiratory disorders or recent (within 3 months) self-reported respiratory tract infection or antibiotic use were excluded. Approval was obtained from the Local Research Ethics Committee (Ref 00/BA/459E and 07/H0712/138) and written informed consent was obtained from all subjects.

### Bronchoscopy

Fibre-optic bronchoscopy with BAL was performed via the oropharyngeal route with topical anaesthesia and midazolam sedation in accordance with American Thoracic Society guidelines.^16^ Aliquots of buffered warm saline, to a total volume of 240ml, were separately instilled in to a segment of the right middle lobe and gently aspirated by manual suction into a sterile syringe. All return samples were pooled into a 200 ml falcon tube. BAL was undertaken prior to visual inspection of the bronchial tree and in advance of any other planned bronchoscopic procedures. Negative control samples were collected by aspirating buffered saline through the bronchoscope suction channel prior to bronchoscopy. BAL fluid was examined for the presence of macrophages to confirm access of the alveolar compartment, and the absence of ciliated epithelium was used to exclude large airway contamination.

### DNA extraction

Genomic DNA was isolated from BAL using the MP Bio FastDNA^®^ SPIN Kit for Soil (http://www.mpbio.com). Two 2 ml Aliquots of BAL were centrifuged at 20,000 g for 15 minutes, to pellet cell debris and bacteria. Each pellet was then re-suspended in 978 µl Sodium Phosphate Buffer and 122 µl MT Buffer (MPBIO first lysis buffer solution) and extraction performed according to the manufacturer’s protocol. Bead-beating was undertaken using the Precellys 24 bead-beater (PeqLab, UK; 6000 rpm [30 seconds × 2]), as per the manufacturer’s protocol. Isolated DNA was stored at −20°C until use.

### 16S rRNA gene qPCR and Pyrosequencing

Analyses of DNA from controls and IPF samples were performed at the same time by a single operator (PLM). The V3-V5 region of the bacterial 16S rRNA gene was amplified using the 357F forward primer^17^ and the 926R reverse primer^18^ for both 16S qPCR and pyrosequencing. Triplicate 10 µl qPCR reactions were set up containing 1 µl of a 10-fold dilution of template DNA as previously described.^19^ All data were analyzed using the Corbett rotor-gene 6.1 software.

For pyrosequencing the V3-V5 region of the bacterial 16S rRNA gene was amplified using the standard forward primer 357F and a modified reverse primer 926R, barcoded to tag each PCR product. Quadruplicate 25 µl PCR reactions were set up, amplified, purified and prepared for sequencing as previously described.^19^ Single direction pyrosequencing using the Lib-L kit was performed using the 454 Life Sciences GS Junior (Roche). The barcoded pyrosequence reads were processed using QIIME 1.7.0.^20^ Initial denoising was performed to remove sequencing errors^21^ and PCR generated artefacts were removed using ChimeraSlayer.^22^ Sequence reads were removed if they contained ambiguous base calls, mismatches in the primer sequences, homopolymer runs or a mean window quality score less than 25. Sequences were clustered into Operational Taxonomic Units (OTUs) at 97% identity,^23^ aligned to full length 16S rRNA sequences^24^ and assigned a taxonomic identity with the Ribosomal Database Project classifier^25^ using the SILVA reference database.^26^ Any sequences present only once (singletons) were removed.

### Statistical analysis

Continuous variables are presented as means (±Standard Deviation [SD]), and categorical variables as proportions. Metastats was used to perform non-parametric *t*-test comparisons of microbial communities between groups,^27^ with *P*-values corrected by multiple hypotheses testing using the FDR approach of Benjamini and Hochberg. We restricted testing to OTUs that had a differing mean abundance between cases and controls of more than 1% of the total. Shannon’s entropy^28^ (alpha diversity index) and weighted and unweighted UniFrac distances^29^ (beta diversity) were calculated in QIIME. The time-to-event curves were calculated using the Kaplan–Meier method and compared with the use of the log-rank test. Differences between subject groups were evaluated with the use of the Mann–Whitney test for continuous variables and Fisher’s exact test for categorical variables. Pearson’s correlation was used to calculate correlations between continuous variables. The statistical significance of association of variables with a diagnosis of IPF was assessed with the use of stepwise logistic regression. All analyses were performed with the use of Systat 13 and R (http://cran.r-project.org/). A two-sided *P* value of less than 0.05 was considered to indicate statistical significance.

## Results

Seventy five patients with suspected IPF and 44 controls (27 healthy smokers and non-smokers and 17 subjects with moderate COPD (Supplementary Table 1)) were enrolled in the study and underwent bronchoscopy according to the same protocols. During the study period, nine patients with suspected IPF were not enrolled as they reported symptoms of a LRTI or antibiotic usage in the preceding three months. Ten of the 75 recruited patients were excluded because they did not fulfil the ATS diagnostic criteria for IPF. The remaining 65 IPF subjects were predominantly male (77%) with a mean age of 68 years, and had moderately severe disease at enrolment (Carbon Monoxide Diffusing Capacity [DLco] 44.7% predicted; forced vital capacity [FVC] 76.5% predicted). The controls were matched for smoking history and sex but were slightly younger than the IPF cohort with a mean age of 58.2 years (Table 1). None of the control group and only four of the IPF subjects were using inhaled corticosteroids at the time of their assessment.

**Table 1.** Baseline Characteristics of the Patients

| Characteristics | IPF<br>(n=65) | Controls<br>(n=44) | P<br>Value |
| --- | --- | --- | --- |
| Age (yr) | 68 (±8.2) | 58.2 (±8.0) | < 0.0001 |
| Female Sex - number (%) | 15 (23%) | 17 (28%) | ns |
| Smoking (Ever vs Never) - number (%) | 43 (65%) | 28 (63%) | ns |
| Forced vital capacity (FVC)— % | 76.5 (± 18) | 100 (±14) | < 0.0001 |
| Forced Expiratory Volume in 1 Second (FEV<sub>1</sub>) — % | 77 (±15) | 88.5(±19) | < 0.005 |
| Ratio of FEV<sub>1</sub> to FVC | 71.3 (±11) | 80.0 (±7) | < 0.0001 |
| Carbon monoxide diffusion capacity - % of predicted value | 44.7 (±13) | 78.8 (±18) | < 0.0001 |
| O<sub>2</sub> Saturation— % | 94.8 (±2) | 96.9 (±4) | < 0.0001 |
| 6-Minute walk distance - m | 377.8 (±124) | NA | NA |
| Inhaled Steroid Usage | 4 | 0 | ns |
| Bronchoalveolarlavage culture Positive | 5 (7.5%) | 0 | ns |

Data are means ±Standard Deviation

Genomic DNA was successfully extracted from all 109 subjects. Following denoising, chimera checking and singleton removal a total of 912,883 high quality 16S rRNA reads remained. The distribution of reads was between 796 and 12,329 per sample. In order to control for bias of per sample read coverage, sequences were randomly resampled (rarefied) to the same minimum of 796 reads for all subjects. The bacterial sequences were then clustered by sequence similarity into OTUs and this final dataset was used in all following analyses. There were 464 OTUs identified across the IPF and control panels and less than 5% of the sequences were unclassifiable by reference to the SILVA reference database.

IPF subjects had on average 3.9×10^9^ copies of the 16S rRNA gene per ml of BAL, which was more than two-fold higher than the copy number in controls (*P*<0.0001) (Figure 1). Within the controls, there was no significant difference in bacterial load between the subjects with COPD and the healthy controls, while the IPF subjects had a significantly higher bacterial burden than both individual control subgroups (*P=*0.006 and *P*=0.0007 respectively) (Figure 1). Within the IPF subjects there was no correlation between bacterial burden and baseline disease severity, measured by either FVC (Spearman’s ρ=-0.11, P=0.49), total lung capacity (TLC) (Spearman’s ρ=-0.12, *P*=0.49), carbon monoxide diffusion capacity (DLCO) (Spearman’s ρ=-0.20, *P*=0.21) or the composite physiologic index (CPI) (Spearman’s ρ=0.06, P=0.70). Although BAL yield differed between IPF cases and controls (124.4mls ± 31 vs. 96mls ± 41; *P*<0.001) there was no relationship between bacterial burden and BAL yield (Spearman’s ρ=-0.027, *P*=0.78). No ciliated epithelial cells (that may have been indicative of large airway contamination) were seen in the BAL returns. The negative control samples yielded a bacterial burden close to or below the lower limit of qPCR quantification (1000 copies per ml).

**Figure 1:**
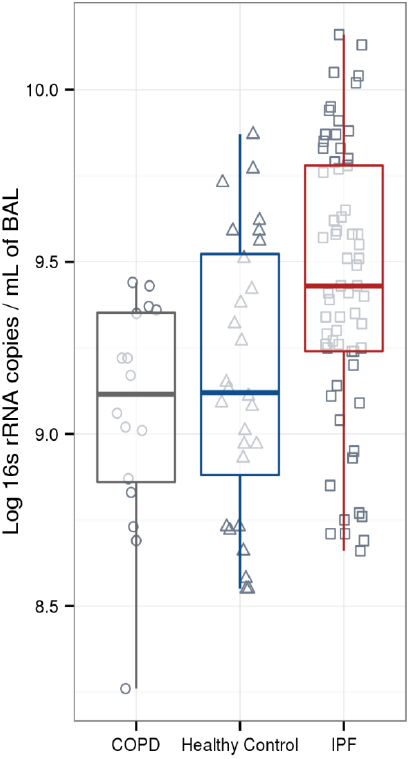
Bacterial Burden in IPF compared to controls. IPF patients (red) had a significantly higher bacterial burden than both subjects with COPD (grey) and the healthy controls (blue) (P=0.006 and P=0.0007 respectively [Mann–Whitney test]). Triangles represent COPD subjects, circles healthy controls and squares IPF subjects. The box signifies the 25th and 75th percentiles, and the median is represented by a short black line within the box.

Although the course of IPF is variable, a 6 month decline of greater than 10% in FVC is associated with an increased risk of mortality.^30^ We separated the patients into subjects with progressive or stable disease by defining disease progression as a relative decline in FVC of greater than 10% or death. Individuals with IPF whose disease had progressed at six months (n=22) demonstrated a significantly higher BAL bacterial burden when compared to subjects with stable disease (4.70×10^9^ ±7.20×10^8^ vs. 2.82×10^9^ ±5.08×10^8^ copies of the 16S rRNA gene per ml of BAL; *P*=0.02). In order to investigate this further the IPF subjects were separated into tertiles based on the 16S rRNA gene copy number per ml of BAL. Individuals in the top tertile with the highest bacterial burden were at substantially increased risk of mortality compared to subjects in the bottom tertile with the lowest bacterial burden (hazard ratio 4.59 (95% Confidence Interval [CI], 1.05-20) (Figure 2).

**Figure 2:**
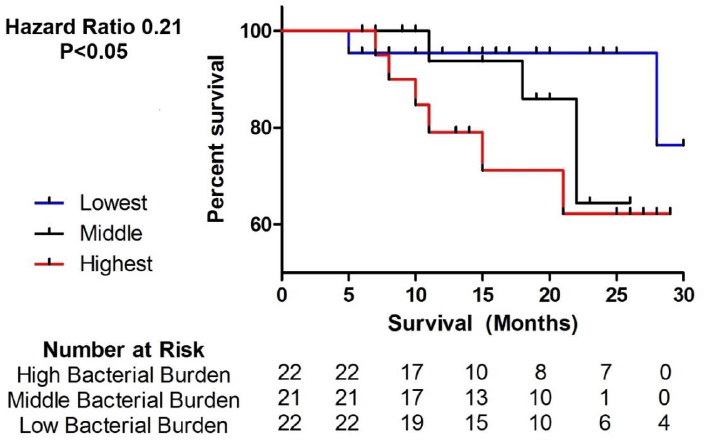
Kaplan–Meier Curves for the Time until Death. Subjects with IPF in the tertile with the highest bacterial load (16S copy number per mL of bronchoalveolarlavage) are shown in red, and were at substantially increased risk of mortality compared to IPF subjects in the tertile with the lowest bacterial burden, shown in blue.

Standard microbial culture of BAL was positive in 5 cases of IPF (7.6%) but none of the controls. In each instance, the cultured bacterial species were also identified by the pyrosequencing data, although they were not always the most abundant species within the microbiome.

*Streptococcus,* representing 30% of total reads, was the most common genus detected by sequencing in subjects with IPF. *Prevotella* (10.9%) and *Veillonella* (10.6%) were the second and third most prevalent. In the control subjects this order was preserved with *Streptococcus* forming the most common genus (27.1% of total reads) followed by *Prevotella* (11.6%) and *Veillonella* (7.1%). The microbial communities of IPF subjects were less diverse (Shannon’s diversity index 3.81 ± 0.08 vs. 4.11 ± 0.10; *P*=0.005) and contained fewer numbers of OTUs (44.89 ± 1.50 vs. 54.33 ± 1.86 OTUs; *P*< 0.0001) than the control subjects. OTUs that were present only in control subjects were well below 1% relative abundance, so the biological significance of the difference in diversity is uncertain.

Specific differences were observed between cases and controls at the phylogenetic level (Supplementary Figure 1). Four OTUs showed an increased number of reads in the IPF cohort compared to controls (Figure 3). A 3.4 fold increase was observed in sequence numbers of a potentially pathogenic *Haemophilus* sp. (OTU 739) (36.0 ± 7.5 vs. 10.5 sequences ±1.9 sequences; *P*<0.001); a 2.1 fold increase in a *Neisseria* sp. (OTU 594) (57.9 ± 9.4 vs. 27.5 ± 5.5 sequences; *P*<0.01); a 1.4 fold increase in a *Streptococcus* sp. (OTU 881) (113.6 ± 11.4 vs. 82.2 ± 8.9 sequences; *P*<0.05); and a 1.5 fold increase in a *Veillonella* sp. (OTU 271) (84.8 ± 5.7 vs. 56.6 ± 4.5 sequences; *P*<0.001). Despite the differences in overall bacterial burden there were no demonstrable changes in community structure or composition between IPF patients with declining or stable lung function. There were no significant differences in the BAL microbiota between the healthy controls and subjects with COPD, and exclusion of subjects with COPD from the control panel did not change the differences detected.

**Figure 3:**
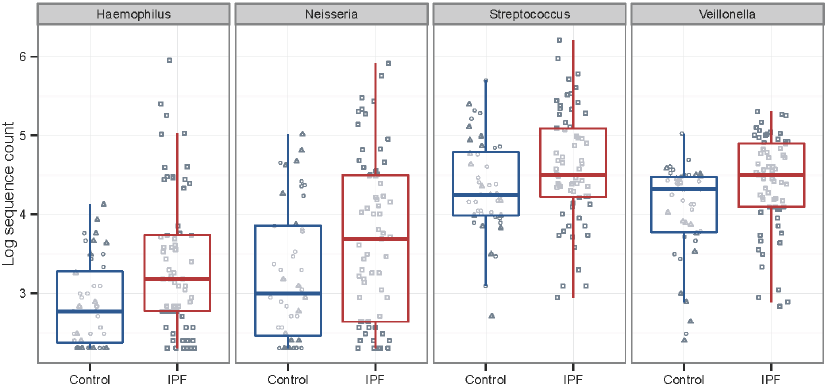
Differences in Bacterial OTU frequencies between IPF and control subjects. Box plots showing significant differences (*P*<0.01) in OTUs between controls (blue) and patients with IPF (red). Triangles represent COPD subjects, circles healthy controls and squares IPF subjects. The box signifies the 25th and 75th percentiles, and the median is represented by a short black line within the box.

A stepwise logistic regression was undertaken to determine if bacterial burden and the relative abundance of these bacteria associated independently with a diagnosis of IPF. Age was included in the model to control for possible confounding. Total bacterial burden (log copy number) and the numbers of three specific OTUs (*Veillonella* sp. [OTU 271], *Neisseria* sp. [OTU 594] and *Streptococcus* sp. [OTU 881]) all remained significantly associated with a diagnosis of IPF (Table 2). The overall R^2^ was 0.66 with age in the model and was 0.51 without age. These data indicate that total bacterial load and abundances of three specific OTUs provide independent predictors of IPF case status.

**Table 2.** Stepwise logistic regression of bacterial associations to Idiopathic Pulmonary Fibrosis compared to controls

|  | Beta | SE | Z | P | 95% CI |  |
| --- | --- | --- | --- | --- | --- | --- |
| **Constant** | -34.41 | 8.740 | -3.938 | <0.001 | -17.28 | -51.54 |
| **Age** | 0.173 | 0.041 | 4.255 | <0.001 | 0.253 | 0.094 |
| ***Veillonella* (OTU 271)** | 0.032 | 0.009 | 3.528 | <0.001 | 0.050 | 0.014 |
| ***Streptococcus* (OTU 881)** | 0.012 | 0.004 | 2.712 | 0.007 | 0.018 | 0.003 |
| **Log<sub>10</sub> Copy Number** | 2.140 | 0.824 | 2.596 | 0.009 | 3.755 | 0.524 |
| ***Neisseria* (OTU 594)** | 0.020 | 0.008 | 2.574 | 0.010 | 0.035 | 0.005 |

Naglekerke's $R^2=0.66$ . Not in the equation: *Haemophilus* sp. (OTU 739), $P=0.25$

The abundance of *Haemophilus* OTU 739 was strongly correlated with *Neisseria* OTU 594 (Spearman’s ρ=0.42): removal of *Neisseria* from the model made *Haemophilus* OTU 739 significant (*P*=0.02; β=0.038±0.016) with minimal change in the overall multivariate R^2^ (0.657). The high β (effect size) relative to other OTUs (Table 2) suggests that *Haemophilus* should not be excluded as a potential source of individual impact on the presence of IPF.

## Discussion

In this study we have shown that when compared to controls, patients with IPF have a higher bacterial load in bronchoalveolar lavage fluid and significant differences in the composition and diversity of their microbiota. We have also shown that an increased bacterial load at the time of diagnosis identifies patients with more rapidly progressive IPF and a higher risk of mortality.

The baseline bacterial communities we observed in IPF patients and control subjects contained organisms such as *Streptococcus, Prevotella, Fusobacterium,* and *Haemophilus* which are commonly found in the airways of healthy subjects, asthmatics and COPD patients.^31–34^ We discovered differences in specific OTUs between cases and controls, notably the presence of more *Streptococcus*, *Haemophilus*, *Neisseria* and *Veillonella* spp. in the IPF patients.

Our results suggest that OTUs representing potential pathogens such as *Haemophilus* or *Streptococcus* may be acting synergistically within a context of increased load. The lack of an association between baseline disease severity and bacterial load suggests that bacteria in the lungs of IPF patients are not a simple consequence of disease extent but that they may in some way contribute to disease progression.

The more densely populated and less diverse microbiota in IPF subjects may provide a low level antigenic stimulus for repetitive alveolar injury. The temporal and spatial heterogeneity observed in usual interstitial pneumonia (the histological lesion of IPF) speaks to the likely importance of repetitive injury as a major factor in the pathogenesis of the disease.^35^ The bacterial communities of the lower airways are a plausible candidate for this trigger, and local differences in the bacterial microbiome^36^ may contribute to the recognised geographical differences in the incidence of IPF.

The increased bacterial load and altered bacterial communities we have discovered in our patients could reflect decreased clearance or impaired immune responses, consistent with the observations that genetic variants in mucin and surfactant genes predisposes to the disease.^4,8^ It will be of particular interest to test if these known genetic variants correlate with characteristics of the microbial community in BAL.

Our study has a number of limitations. IPF is a disease of the lung parenchyma whereas we utilized BAL fluid to sample the distal airways. Direct sequencing of lung tissue may provide further information on the pulmonary microbiome,^37^ but lung biopsies are undertaken relatively infrequently in IPF and gathering true healthy control samples would be difficult. BAL is part of usual clinical practice in patients with IPF, and provides effective accesses to the alveolar space with much less morbidity than biopsy.

We used the oropharnygeal route to pass a bronchcoscope into the lungs, and some secretions from the upper airways will have been carried on the tip of the bronchoscope to the site of wedging for lavage in the right middle lobe. Any carry over will have been heavily diluted by the 240ml of BAL fluid. The high percentage of BAL return and an absence of ciliated epithelial cells provide confidence that our BAL return was primarily derived from the distal airspace. Most importantly, we used the same protocols and procedures for cases and controls, so any contamination will not have systematically influenced our results.

We note that strong similarities have been seen between microbiota identified from BAL fluid and from surgically removed lung tissue in which upper airway contamination was not an issue.^36^ Techniques such as antimicrobial mouthwashes before bronchoscopy^38,39^ and the use of different bronchoscopes for supraglotic anaesthesia and exploration of the lower airway^40^ deserve further validation for use with culture-independent microbial investigations.

Some bacterial species may not have been detected by our PCR primers, and we did not assess for the presence or absence of other potential non-bacterial lung pathogens. Although a substantial study found no evidence of detectable viruses by PCR of BAL fluid from 40 stable IPF patients^11^ it is conceivable that some differences in progression could be due to the presence of respiratory viruses in our sample.

Overt respiratory infection in patients suffering from IPF is known to carry a high morbidity and mortality,^12^ and immunosuppression leads to poor clinical outcomes.^13^ We have demonstrated that an increased bacterial load at the time of diagnosis of IPF is associated with increased mortality. It is not possible to conclude whether the presence of a higher bacterial load and an altered microbiome is the cause or a result of the destruction of the normal lung architecture in IPF. It may be helpful to explore any changes in bacterial burden that precede or accompany acute exacerbations of the disease. One other means to establish causality may be the targeting of specific bacteria with antibiotic therapies in an attempt to reduce mortality in this otherwise intractable disease.

## Acknowledgements

PLM was an Asmarley Clinical Research Fellow. Funding was from the Asmarley Trust, the Wellcome Trust, Medical Research Council Programme Grant G0600879. The project was supported by the NIHR Respiratory Disease Biomedical Research Unit at the Royal Brompton and Harefield NHS Foundation Trust and the AHSC Biomedical Research Centre at Imperial College London. Control samples were collected with an unrestricted grant from GlaxoSmithKline and a grant from Pfizer UK. The late Dr Joseph Footitt was instrumental in the study of controls.

## Author contributions

PLM planned the project, recruited subjects, designed and performed experiments, analysed the data and wrote the manuscript; MJC and SAGWO performed statistical analyses, interpreted the results and helped write the manuscript; KER performed experiments and interpreted the results; PM, JF, SLJ, AMR and AUW enrolled subjects and performed the research; MFM and WOC conceived the microbiological studies of IPF; TMM conceived the systematic prospective study of IPF patients; MFM, WOC and TMM planned the project, designed experiments, analysed the data and wrote the manuscript. All authors reviewed, revised and approved the manuscript for submission.

---

**Supplementary Table 1:**
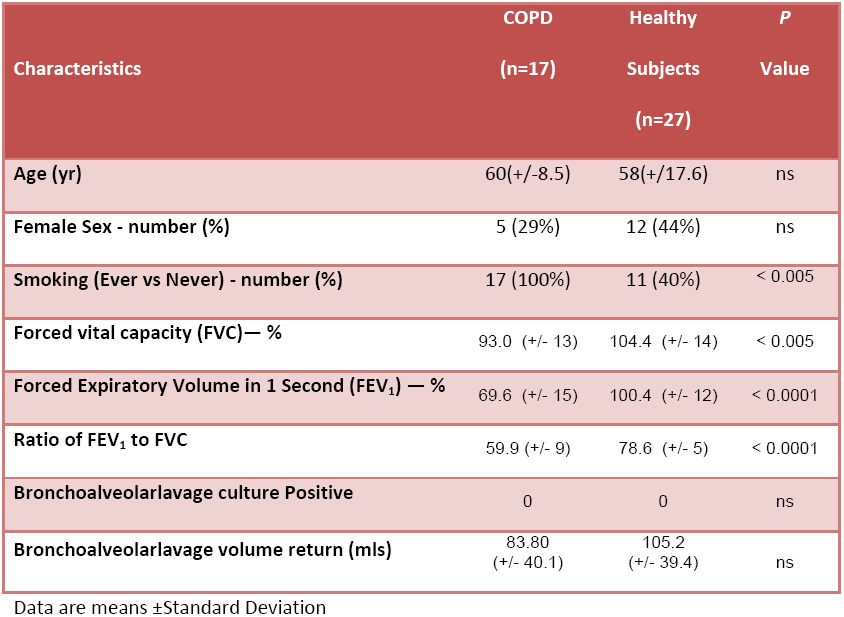
Baseline Characteristics of the Control patients split into healthy individuals and COPD subjects.

| Characteristics | COPD<br>(n=17) | Healthy<br>Subjects<br>(n=27) | P<br>Value |
| --- | --- | --- | --- |
| Age (yr) | 60(+/-8.5) | 58(+/-17.6) | ns |
| Female Sex - number (%) | 5 (29%) | 12 (44%) | ns |
| Smoking (Ever vs Never) - number (%) | 17 (100%) | 11 (40%) | < 0.005 |
| Forced vital capacity (FVC)— % | 93.0 (+/- 13) | 104.4 (+/- 14) | < 0.005 |
| Forced Expiratory Volume in 1 Second (FEV<sub>1</sub>) — % | 69.6 (+/- 15) | 100.4 (+/- 12) | < 0.0001 |
| Ratio of FEV<sub>1</sub> to FVC | 59.9 (+/- 9) | 78.6 (+/- 5) | < 0.0001 |
| Bronchoalveolarlavage culture Positive | 0 | 0 | ns |
| Bronchoalveolarlavage volume return (mls) | 83.80<br>(+/- 40.1) | 105.2<br>(+/- 39.4) | ns |

Data are means ±Standard Deviation

**Supplementary Figure 1.**
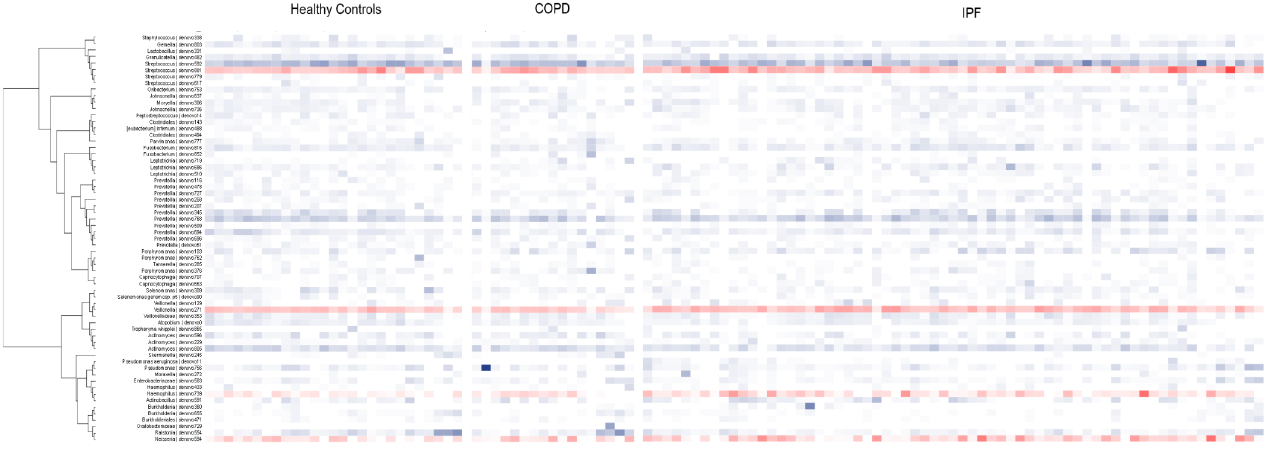
Phylogenetic tree and heatmap of bacterial 16S rRNA sequences grouped into IPF, COPD and healthy controls. This depicts the top 100 OTUs organised phylogenetically by tree with abundance indicated by the colour (Darker Blue or Red more abundant). Taxonomy assignments at the phylum level are shown in the inner column and colour coded and the OTUs of interest are highlighted in red.

